# Avian Influenza A Virus polymerase can utilise human ANP32 proteins to support cRNA but not vRNA synthesis

**DOI:** 10.1101/2022.06.27.497881

**Authors:** Olivia C. Swann, Amalie B. Rasmussen, Thomas P. Peacock, Carol M. Sheppard, Wendy S. Barclay

## Abstract

Host restriction limits the emergence of novel pandemic strains from the Influenza A Virus avian reservoir. For efficient replication in mammalian cells, the avian influenza RNA-dependent RNA polymerase must adapt to use human orthologues of the host factor ANP32, which lack a 33 amino acid insertion relative to avian ANP32A. Here we find that influenza polymerase requires ANP32 proteins to support both steps of replication: cRNA and vRNA synthesis. Nevertheless, avian strains are only restricted in vRNA synthesis in human cells. Therefore, avian polymerase can use human ANP32 orthologues to support cRNA synthesis, without acquiring mammalian adaptations. This implies a fundamental difference in the mechanism by which ANP32 proteins support cRNA vs vRNA synthesis.

**Importance:** In order to infect humans and cause a pandemic, avian influenza must first learn how to use human versions of the proteins the virus hijacks for replication – instead of the avian versions found in bird cells. One such protein is ANP32. Understanding the details of how host proteins such as ANP32 support viral activity may allow the design of new antiviral treatments that disrupt these interactions. In this work, we use cells that lack ANP32 to unambiguously demonstrate ANP32 is needed for both steps of influenza genome replication. Surprisingly however, we find that avian influenza can use human ANP32 proteins for the first step of replication without any adaptation, but only avian ANP32 for the second step of replication. This suggests ANP32 may have an additional role in supporting the second step of replication, and it is this activity that is specifically blocked when avian influenza infects human cells.

## Introduction

Influenza A viruses (IAVs) pose a pandemic risk: whilst the natural hosts of IAV are aquatic birds, the virus is associated with sporadic zoonotic jumps, which may trigger widespread disease in an immunologically naïve population (1). The enormous consequences of such events are illustrated by the historical 1918 ‘Spanish flu’ pandemic, and the ongoing coronavirus pandemic caused by SARS-CoV-2.

Prior to emerging as a pandemic strain, zoonotic IAV must adapt to overcome multiple host range barriers. One important restriction is adaptation of the RNA-dependent RNA polymerase (FluPol) for efficient activity within the mammalian cellular environment (1, 2). FluPol is a heterotrimer consisting of three viral proteins: PB1, PB2 and PA. Each of the eight viral genomic RNAs (vRNAs) of the IAV genome are encapsidated by nucleoproteins (NP) and associate with a FluPol protomer in a viral ribonucleoprotein complex (vRNP). During infection, FluPol drives both transcription and replication of the viral genome (3). Transcription occurs in a primer-dependent manner to produce capped and poly-adenylated positive sense viral mRNAs. Genome replication occurs in a two-step process: first, negative sense vRNA is copied into a full-length positive-sense cRNA intermediate which is packaged into complementary RNPs (cRNPs) by acquiring an encapsidating polymerase and NP. Nascent vRNPs are then produced from cRNPs (3, 4). cRNA and vRNA synthesis differ in *de novo* initiation strategy. Whilst cRNA synthesis uses terminal initiation, vRNA synthesis uses internal initiation followed by primer realignment that is dependent on an additional trans-activating FluPol (5–8).

Structural studies suggest the functional flexibility of FluPol is possible due to rearrangements of peripheral domains outside the catalytic core (3). These depend on the presence or absence and nature of the bound RNA promoter, or upon dimerization with additional polymerase molecules (9–11). FluPol also co-opts host factors to co-ordinate its activity, some of which may stabilise different functional states of the polymerase (10, 12).

ANP32 proteins are a family of proviral host factors that are essential for influenza replication (13, 14). Species differences between ANP32 proteins underly host restriction of avian influenza polymerase in mammalian cells (15). Prototypical avian influenza polymerase bearing a glutamic acid at position 627 of the PB2 protein (hereafter referred to as FluPol 627E) cannot be supported by mammalian ANP32 proteins that lack the 33 amino acid insertion present in avian ANP32A orthologues, and accordingly show restricted replication in human cells. In contrast, FluPol bearing a single amino acid substitution at the PB2 627 position from glutamic acid to lysine (the quintessential mammalian-adapting mutation, FluPol 627K), can utilise mammalian ANP32 proteins that lack this insertion, and such viruses replicate to high titres in human cells (2, 15). Interestingly, although chicken ANP32A (chANP32A) shows enhanced binding to FluPol relative to human ANP32A (huANP32A), that is dependent on the 33 amino acid insertion, this pattern occurs independent of FluPol 627 signature, i.e., species-specific functionality is not dictated by differences in binding affinity (16–18).

Previous studies suggest that ANP32 proteins are specifically required to support vRNA synthesis, and accordingly that host restriction occurs at this step of replication. Sugiyama et al found that purified huANP32A stimulated RNA replication from a short cRNA but not vRNA template in *in vitro* replication assays (19). Moreover, studies comparing infection in mammalian cells with strains bearing either FluPol 627E or 627K found that 627E was specifically restricted in vRNA synthesis, but not cRNP stabilisation (20, 21). Nevertheless, other reports implicate ANP32 in both steps of replication (11, 22). The recent structures of huANP32A and chANP32A in complex with FluPol from Influenza C virus demonstrate that the N terminal Leucine Rich Repeat (LRR) domain of ANP32 bridges a novel asymmetric dimer of two FluPol enzymes. The unstructured C terminal Low Complexity Acidic Region (LCAR) domain of ANP32 remains largely unresolved, apart from additional density sandwiched in a groove formed between the two 627 domains. This dimer was interpreted as a ‘replication complex’, with the RNA-bound FluPol (FluPol_R_) adopting a replication-competent structure, and the additional apo-enzyme (FluPol_E_) adopting a novel conformation: poised to encapsidate the nascent RNA into an RNP complex. Interestingly, this structure was obtained with FluPol_R_ bound to a 47nt short vRNA, i.e., primed for cRNA synthesis, which was not previously implemented in requiring support by ANP32. Moreover, mutations introduced to disrupt the FluPol asymmetric dimer interface resulted in a significant reduction in cRNA encapsidation. This, combined with the logic that both cRNA and vRNA require encapsidation into RNPs during replication, suggests that ANP32 may play a role in cRNA synthesis as well as vRNA synthesis (11).

Here we use ANP32 knockout cell lines to clarify the role of ANP32 proteins in cRNA synthesis. We establish an RNA fluorescence in situ (FISH) assay for directly visualising cRNA and demonstrate that ANP32 proteins are essential for primary cRNA synthesis in authentic infection, as well as under experimental conditions in which vRNA synthesis is inhibited. Nevertheless, we find that avian FluPol 627E does not show restricted cRNA accumulation in mammalian cells. This observation is consistent regardless of whether FluPol_R_, FluPol_E_ or both FluPol molecules in the replication complex bear the avian-like 627E signature. Moreover, the cRNPs produced by FluPol_R_ 627E are functional for onward replication. To conclude, our study suggests that huANP32 proteins are sufficient to support avian FluPol cRNA synthesis, and that host restriction acts specifically at the level of vRNA synthesis. This suggests a fundamental difference in the mechanism by which the host factor ANP32 supports vRNA and cRNA synthesis.

## Results

### cRNA synthesis is inhibited in human cells lacking ANP32A/B

Human cells express three members of the ANP32 protein family: huANP32A, huANP32B and huANP32E (23). Previous work has demonstrated that huANP32A/B are functionally redundant for proviral activity, whilst huANP32E does not support FluPol activity (13, 14). To investigate the role of ANP32 proteins in cRNA synthesis, wild type human eHAP cells (eHAP WT) and human eHAP cells in which ANP32A/B have been ablated (eHAP dKO) (13) were infected with A/Puerto Rico/8/1934(H1N1) (PR8) and the accumulation of segment 4 (haemagglutinin (HA)) vRNA, cRNA and mRNA quantified using a tagged RT-qPCR approach (24). Following infection, there was no increase in either vRNA or cRNA levels over time in the absence of ANP32A/B, confirming ANP32 proteins are essential for replication (Figure 1A,B). Significantly, no increase in cRNA accumulation was observed over input, suggesting a direct role for ANP32 proteins in the pioneering round of cRNA synthesis. In contrast, HA mRNA transcripts increased 50-fold in dKO cells from 0 hours post infection (hpi) to 2hpi, illustrating that primary transcription does not require ANP32 proteins (Figure 1C). In a separate experiment, we analysed the accumulation of segment 6 (neuraminidase (NA)) vRNA, cRNA and mRNA at an early and late timepoint in eHAP WT and dKO cells, using an analogous tagged RT-qPCR approach. As with segment 4, at late timepoints a significant reduction in all three RNA species was observed (Figure 1D,E,F). At 3hpi, a ∼10 fold reduction in the accumulation of cRNA was already apparent in the dKO cells, despite no difference in vRNA accumulation having yet occurred. Again, this suggested a direct role for huANP32A/B in supporting cRNA synthesis.

**Figure 1.**
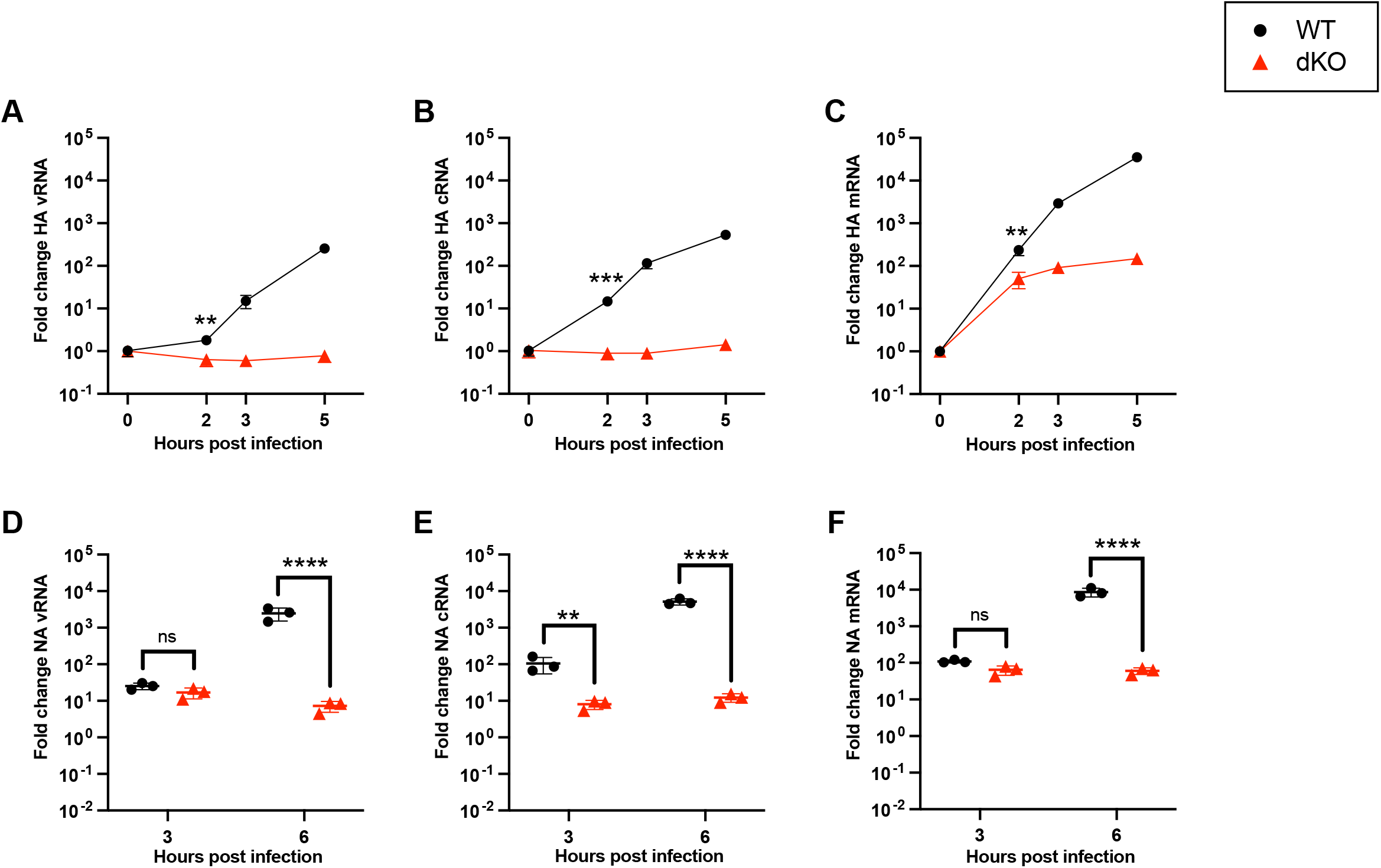
cRNA synthesis is inhibited in human cells lacking ANP32A/B. (A-C) Segment 4 (A) vRNA (B) cRNA or (C) mRNA accumulation over time in eHAP WT vs dKO cells following infection with PR8 (MOI=3). Fold change was calculated over input (0hpi). n=3 biological replicates, plotted as mean ± s.d.. Significance was assessed at 2hpi using an unpaired t-test following log transformation. (D-F) Segment 6 (D) vRNA (E) cRNA or (F) mRNA accumulation over time in eHAP WT vs dKO cells following infection with PR8 (MOI=3). Fold change was calculated over mock infected cells. n=3 biological replicates, plotted as mean ± s.d.. Significance was assessed using multiple unpaired t-tests following log transformation, corrected for multiple comparisons using false discovery rate. hpi=hours post infection. ns=not significant; **=p<0.01; ***=p<0.001; ****=p<0.0001.

### Human ANP32 proteins play a direct role in primary cRNA synthesis

To eliminate the possibility that ANP32 proteins are only required for vRNA synthesis, and the lack of cRNA accumulation is an indirect effect, we made use of cRNP stabilisation assays. cRNP stabilisation assays manipulate cellular conditions to allow only the primary round of cRNA synthesis from incoming viral genomes (Vreede et al, 2004, Nilsson et al, 2017). vRNA synthesis, and consequently further secondary rounds of cRNA synthesis, are inhibited. The assay involves pre-expressing FluPol and NP, and then infecting cells with virus. Importantly, the FluPol that is exogenously expressed is a catalytically dead mutant (PB1 D446Y) that will allow stabilisation of nascent cRNA into catalytically inactive cRNPs. Cells are then drug treated with either actinomycin D (ActD) or cycloheximide (CHX) during infection to inhibit the synthesis of nascent viral proteins which would otherwise enable normal replication.

Initially, ActD-treated cRNP stabilisation assays were performed in eHAP WT and dKO cells. ActD inhibits viral transcription, therefore in these assays only primary cRNA synthesis occurs (Figure 2A). When analysed by RT-qPCR (Figure 2B,C,D) a highly significant reduction in cRNA was observed in dKO cells as compared to WT cells. As expected, no accumulation of NA vRNA or mRNA occurred over background levels (controls lacking transfected PB1, dotted line), in either cell type. Equal transfection efficiency was confirmed by western blot (Figure S1A).

**Figure 2.**
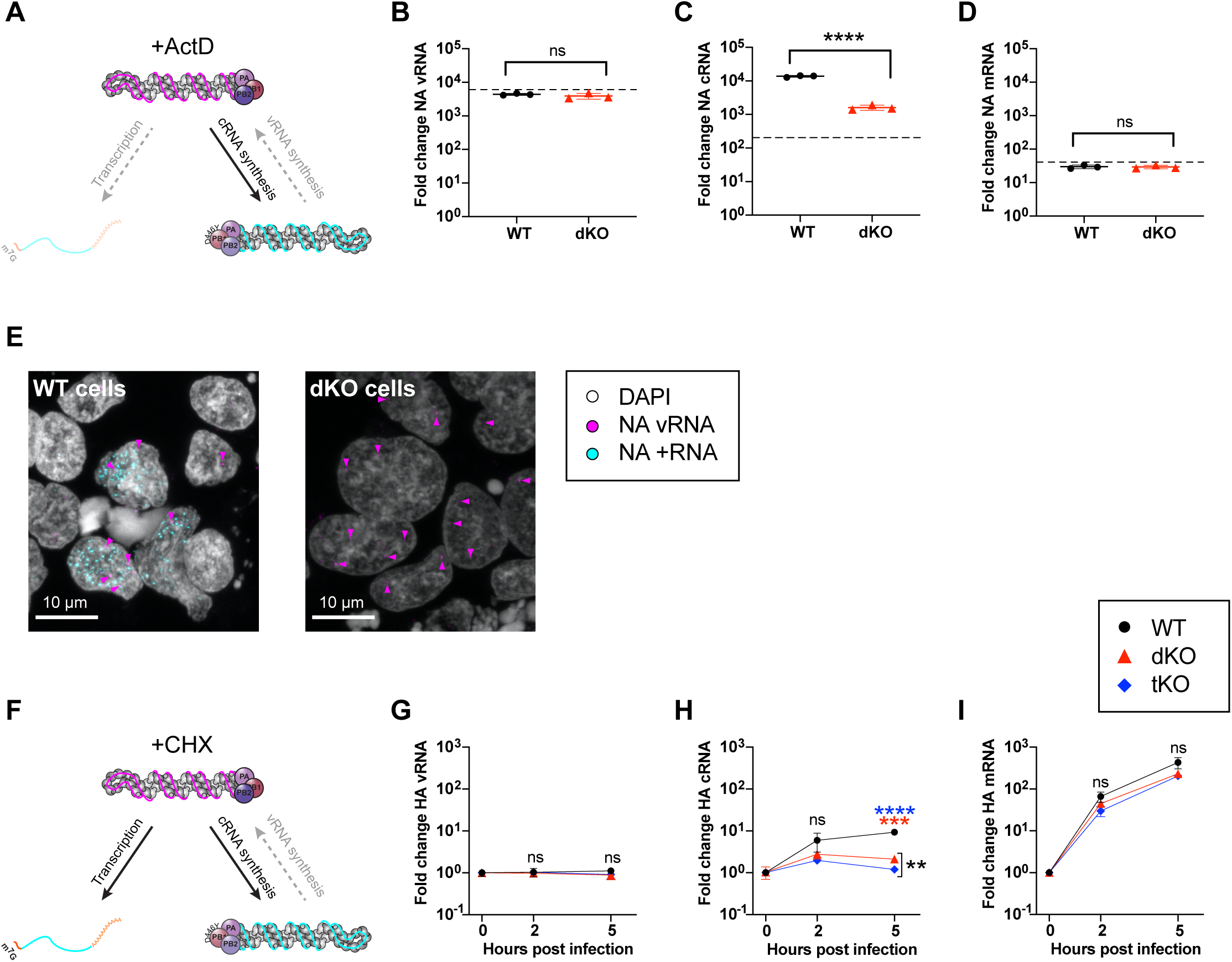
Human ANP32 proteins play a direct role in primary cRNA synthesis. (A) Schematic illustrating influenza polymerase activity under the conditions of a cRNP stabilisation assay with ActD. (B-E) cRNP stabilisation assay with ActD in eHAP WT vs dKO cells, following infection with PR8 (MOI=3), 3hpi. Segment 6 (B) vRNA (C) cRNA or (D) mRNA accumulation. Dotted line indicates background RNA present in control samples transfected with a plasmid mix lacking PB1. Fold change was calculated over mock infected cells. n=3 biological replicates, plotted as mean ± s.d.. Significance was assessed using an unpaired t-test following log transformation. (E) Accumulation of segment 6 vRNA and +RNA in eHAP WT and dKO cells, analysed using RNAscope. Magenta arrowheads highlight a subset of NA vRNA-stained puncta. Images are representative maximum intensity projections. (F) Schematic illustrating influenza polymerase activity under the conditions of a cRNP stabilisation assay with CHX. (G-I) cRNP stabilisation assay with CHX in eHAP WT, dKO and tKO cells. Segment 4 (G) vRNA (H) cRNA or (I) mRNA accumulation following infection with PR8 (MOI=3). Fold change was calculated over input (0hpi). n=3 biological replicates, plotted as mean ± s.d.. Significance was assessed using multiple unpaired t-tests following log transformation, corrected for multiple comparisons using false discovery rate. hpi=hours post infection. ns=not significant; **=p<0.01; ***=p<0.001; ****=p<0.0001.

We next established *in situ* assays for directly visualising IAV replication products to allow single cell, spatial information to be collected alongside bulk assay readouts. To achieve this, the RNA FISH assay RNAscope® was used (25). Probes were designed to target either segment 6 negative sense RNA (NA vRNA probe) or positive sense RNA (NA +RNA probe). The NA +RNA probe is unable to distinguish NA cRNA/mRNA due to the minimal sequence differences between these two RNA species. To counter this, simultaneous cRNP stabilisation and replication assays were performed with ActD treatment, to inhibit viral transcription and allow +RNA staining to be attributed to cRNA. Replication assays are performed as for cRNP stabilisation assays, however active polymerase is pre-expressed, so that multiple rounds of cRNA and vRNA synthesis take place. Validation of the approach is shown in Figure S1B-F. In an *in situ* cRNP stabilisation assay, +RNA accumulation could be clearly detected 3hpi in the nuclei of infected WT cells (Figure 2E). In contrast, whilst incoming vRNPs could be observed in the nuclei of dKO cells, no +RNA staining was observed. This corroborates the view that huANP32A/B are required for primary cRNA synthesis.

Our infection data (Figure 1) suggest that ANP32 proteins are not required for primary transcription. Nevertheless, at later infection timepoints, reduced mRNA accumulation is observed in dKO cells, compared to WT cells (Figure 1C,F). To confirm this is an indirect effect due to reduced vRNP template, we chose to undertake a CHX-treated cRNP stabilisation assay in eHAP wild type, dKO or in eHAP cells lacking expression of all three huANP32 proteins: A, B and E (eHAP tKO). In this version of the assay, both cRNA synthesis and primary transcription occur in the absence of vRNA synthesis (Figure 2F). No accumulation of vRNA over input was observed in any of the cell types over time (Figure 2G), as expected. In agreement with ActD cRNP stabilisation assays (Figure 2C), a significant decrease in cRNA accumulation was seen in both dKO and tKO cells compared to WT by 5hpi (Figure 2H). A small but significant difference (p=0.0018) was observed between cRNA accumulation in the dKO and tKO cells, implying huANP32E may support a low level of cRNP stabilisation. In contrast, no significant difference was observed in mRNA accumulation in the presence or absence of any ANP32 proteins from 0 to 5hpi (Figure 2I). This confirms that primary transcription does not require ANP32 proteins. Comparable transfection efficiency was confirmed via western blot (Figure S2G).

### Avian polymerase is not restricted in cRNA synthesis in mammalian cells

Restriction of avian signature FluPol (FluPol 627E) in mammalian cells is attributed to species differences in ANP32 proteins. Consequently, as we have confirmed that ANP32 is required to support both cRNA and vRNA synthesis, we would expect FluPol 627E to be restricted in both steps of replication. Nevertheless, previous work maps restriction to occur specifically at the level of vRNA synthesis (20, 21, 26). To investigate this apparent contradiction, we used a pair of isogenic viruses based on the avian strain A/turkey/England/50-92/1991 (H5N1) (5092) that differ only in the residue at PB2 position 627: either the wild type PB2 627E (hereafter referred to as 5092E) or the humanising mutation PB2 E627K (5092K) (27).

First, we confirmed that 5092 polymerase is also dependent on ANP32 proteins for cRNA synthesis both during authentic infection (Figure S2A,B,C) and in cRNP stabilisation assays (Figure S2D,E,F). Next, we undertook simultaneous cRNP stabilisation and replication assays in human cells, where only the shorter huANP32A/B/E proteins, that are incompatible with FluPol 627E, are available. We pre-expressed NP with either catalytically dead 5092 FluPol 627E or 627K (cRNP stabilisation assay) or WT 5092 FluPol 627E or 627K (replication assay), followed by infection with either 5092K or 5092E virus in the presence of ActD. Significantly, these assays allow the effect of PB2 residue 627 in either FluPol_R_ or FluPol_E_ to be differentiated. The resulting FluPol combinations for both cRNA and vRNA synthesis are outlined in Figure 3A.

**Figure 3.**
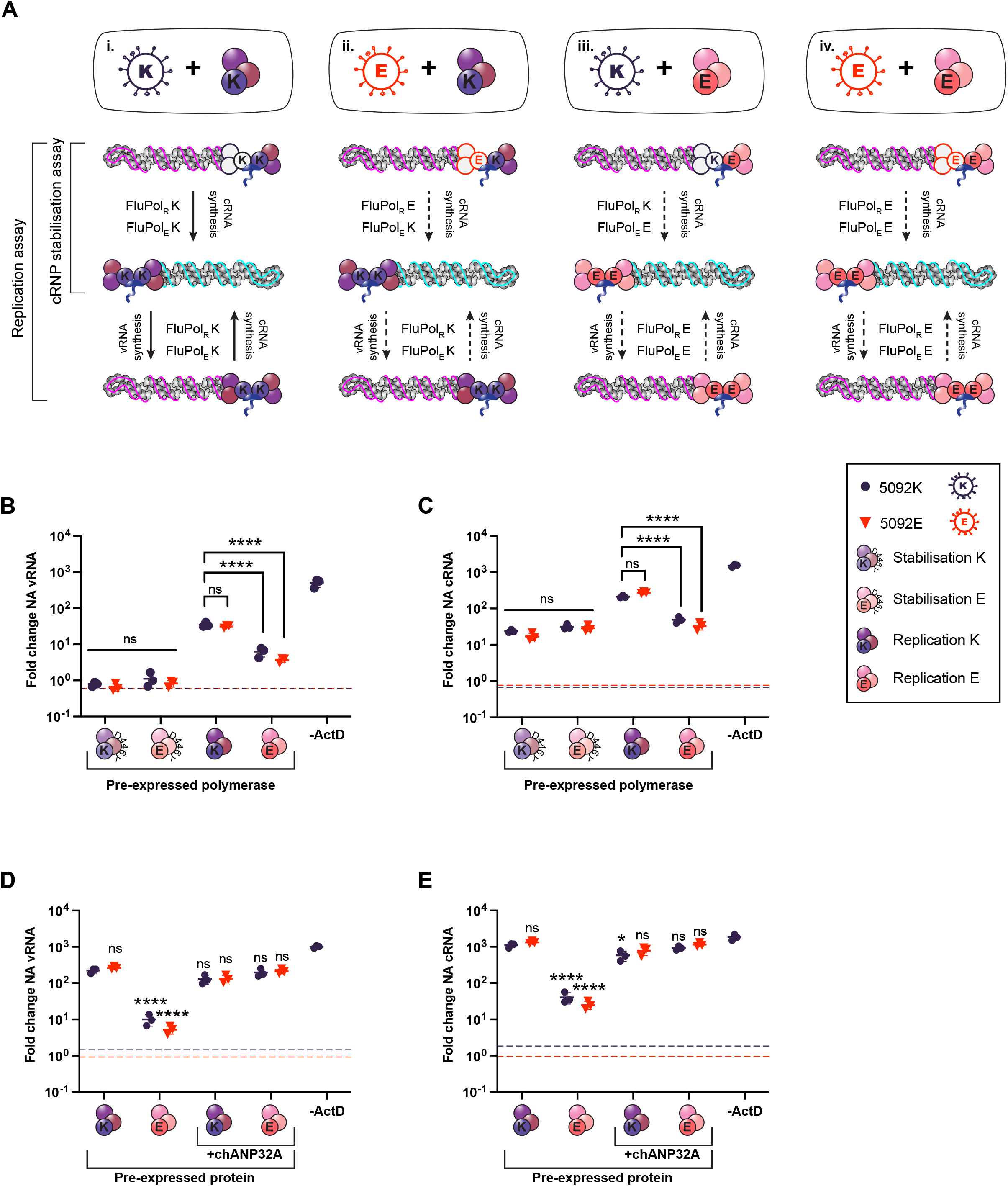
Avian polymerase is not restricted in cRNA synthesis in mammalian cells. (A) Schematic outlining assay set-up and expected polymerase combinations during replication. As indicated on the left, in the cRNP stabilisation assay, only the first layer of activity (primary cRNA synthesis) will occur. Both layers of activity can occur in replication assays. Unknown polymerase activity is indicated by a dashed black arrow. (B,C) Simultaneous cRNP stabilisation and replication assays with ActD, 6 hpi. Segment 6 (B) vRNA and (C) cRNA accumulation following infection with either 5092E or 5092K as indicated (MOI=0.1). Dotted line indicates levels of vRNA/cRNA present in a control lacking PB2 in the transfection mix. Fold change was calculated over input (0hpi). n=3 biological replicates, plotted as mean ± s.d.. Significance was assessed using one way ANOVA with Dunnett’s multiple comparison test, following log transformation. (D,E) Replication assays with ActD and chANP32A, 6 hpi. Segment 6 (D) vRNA and (E) cRNA accumulation following infection with either 5092E or 5092K as indicated (MOI=0.1). Dotted line indicates levels of vRNA/cRNA present in a control lacking PB2 in the transfection mix. Fold change was calculated over input (0hpi). n=3 biological replicates, plotted as mean ± s.d.. Significance was assessed using one way ANOVA with Dunnett’s multiple comparison test, following log transformation. Pre-expressed polymerase mixes: Stabilisation K=PB2 627K/PB1 D446Y/PA/NP; Stabilisation E=PB2 627E/PB1 D446Y/PA/NP; Replication K=PB2 627K/PB1/PA/NP; Replication E=PB2 627E/PB1/PA/NP. hpi=hours post infection. ns=not significant; *= p<0.05; ****=p<0.0001.

During the pioneering round of cRNA synthesis, measured in the cRNP stabilisation assay, FluPol_E_ is provided by the pre-expressed polymerase whilst FluPol_R_ is brought in on the vRNPs from the infecting virus. No vRNA accumulated in the cRNP stabilisation assay, as expected (Figure 3B). Interestingly, neither the PB2 signature of FluPol_R_ nor FluPol_E_ impacted cRNP stabilisation, with all four FluPol combinations accumulating equivalent amounts of cRNA (Figure 3C). This demonstrates that avian and human signature virus can undergo cRNA synthesis equally well in human cells, i.e., avian FluPol is compatible with huANP32 proteins for cRNA synthesis and stabilisation.

In the replication assay, samples with pre-expressed FluPol 627K accumulated equivalent quantities of vRNA and cRNA, regardless of whether the incoming virus was 5092K or 5092E (Figure 3Ai,ii,B,C). Similarly, samples with pre-expressed FluPol 627E were equally impaired in vRNA and cRNA synthesis, independent of infecting virus signature (Figure 3Aiii,iv,B,C). It has previously been described that host restricted FluPol is impaired in nuclear import, due to incompatibilities with the importin-alpha isoforms present in human cells (28). However, this cannot explain the data obtained here, as FluPol 627E was fully functional in supporting the pioneering round of cRNA synthesis and stabilisation within the nucleus during cRNP stabilisation assays. As replication assays allow multiple cycles of replication, the reduced cRNA synthesis observed with pre-expressed FluPol 627E is likely due to the secondary effect of reduced vRNP template. Exogenous expression of chANP32A was able to rescue replication for all FluPol combinations (Figure 3D,E).

## Discussion

Here we have used cell lines that lack expression of ANP32 proteins to unambiguously confirm huANP32 is required to support cRNA synthesis. Nevertheless, we did not observe restriction of avian virus cRNA synthesis in human cells, confirming that host range restriction imparted by species differences in ANP32 acts at the level of vRNA synthesis.

In the structure of the ANP32 and FluPol complex described by Carrique et al, the ANP32 LRR domain bridges the FluPol dimer. The flexible LCAR remains largely unresolved, although there is additional density present in a groove formed between the two FluPol 627 domains. As the groove is acidic in FluPol 627E, the acidic huANP32 LCAR is likely incompatible with this interaction, whereas the mixture of acidic and basic residues in the 33 amino acid insertion of avian chANP32A overcomes this block (11). Our data suggests this mismatch between the huANP32 LCAR and FluPol 627E can be tolerated for cRNA synthesis. This conclusion aligns with the observation that addition of chANP32A to human cells does not stimulate FluPol 627E cRNA synthesis (26).

The ANP32-stabilised replication complex is thought to be required to load an encapsidating FluPol onto the promoter of the extruded strand of nascent c/vRNA. Moreover, it has been proposed that the unstructured ANP32 LCAR domain may act as a molecular whip, which recruits NP for loading onto the nascent RNA (11, 29). Previous studies have suggested that host restricted avian FluPol can generate cRNA in human cells, but that the resulting cRNPs are non-functional for onwards replication (20, 22, 26). However, here we show that cRNA produced from FluPol_R_ 627E is fully functional for vRNA synthesis, provided the encapsidating FluPol bears 627K for onward vRNA synthesis. Therefore, functional cRNPs are produced in the absence of a compatible 627-LCAR interaction, suggesting host restriction does not act at the level of cRNA encapsidation. In combination, our data therefore suggests that the 627-LCAR interaction is required for a further mechanistic role that is specific to vRNA synthesis. For example, a compatible 627-LCAR interaction could be of critical importance to recruit the trans-activating FluPol (York et al., 2013).

In summary, this work has established that huANP32 is required for cRNA synthesis, as well as vRNA synthesis, in human cells. Nonetheless, host restriction of avian FluPol 627E acts specifically at the level of vRNA synthesis, as huANP32 paralogues are sufficient for supporting avian cRNA synthesis and encapsidation. This suggests a fundamental difference in the mechanism by which ANP32 proteins support cRNA vs vRNA synthesis. We have also established an RNA FISH assay which allows visualisation of FluPol replication products. This allows single-cell, spatial information to be collated, which in future could be used to improve our understanding of the spatial elements of FluPol regulation.

## Materials and methods

### Cells and cell culture

Human-engineered Haploid cells (eHAP, Horizon Discovery), eHAP cells with both huANP32A and huANP32B (eHAP dKO) ablated via CRISPR-Cas9, as previously described (13), or eHAP cells with huANP32A, huANP32B and huANP32E ablated via CRISPR-Cas9 (eHAP tKO) (gift from Ecco Staller and Ervin Fodor) were maintained in Iscove’s modified Dulbecco’s medium (IMDM, Thermo Fisher) supplemented with 10% fetal bovine serum (FBS, Labtech), 1% penicillin-streptomycin (pen-strep, Gibco) and 1% non-essential amino acids (NEAA, Gibco). Human embryonic kidney (293Ts, ATCC) and Madin-Darby canine kidney (MDCK, ATCC) cells were maintained in Dulbecco’s modified Eagle medium (DMEM) supplemented with 10% FBS, 1% pen-strep and 1% NEAA. When used for infection, 293T cells were cultured on poly-L-lysine coated plates to aid adherence. All cells were maintained at 37°C and 5% CO_2_.

### Plasmids

pCAGGS expression plasmids encoding the polymerase subunits (PB2, PB1, PA) and NP from PR8 were subcloned from pPolI plasmids. pCAGGS expression plasmids encoding 50-92 PB2, PB1, PA and NP have previously been described (30). The catalytic mutant PB1 D446Y has previously been described (31, 32). pCAGGS expression plasmid encoding FLAG-tagged chicken ANP32 has been described previously (15).

### Viral infections

The full strain names of viruses used in this study are A/Puerto Rico/8/1934(H1N1) (PR8) and A/turkey/England/50-92/1991(H5N1) (50-92). For experiments using 50-92, all infections were performed with recombinant viruses containing the HA, NA and M segments from PR8, the PB1, PA, NP and NS segments from 50-92 and either the WT 50-92 PB2 segment containing a lysine at position 627 (5092E) or a modified PB2 with a glutamic acid at position 627 (5092K), as previously described (27). For infections, virus was diluted in serum-free media to the correct multiplicity of infection (described in figure legends). For comparative 50-92E/K experiments, viral inputs were normalised based on genome copy number. To synchronise infection, viral inoculation was performed at 4°C. In brief, cells were pre-incubated at 4°C for 15 minutes, before addition of viral inoculum and a further incubation at 4°C for 45 minutes. Viral inoculum was then replaced with pre-warmed full media and infected plates incubated at 37°C, 5% CO_2_. At the appropriate time point, cells were processed for RT-qPCR analysis, imaging analysis or immunoblot as described below.

### Replication/cRNP stabilisation assays

For replication/cRNP stabilisation assays, cells were transfected using Lipofectamine 3000 (Invitrogen) with pCAGGS expression plasmid mixtures encoding polymerase components in the ratios 2:2:1:4, PB2:PB1:PA:NP, where 1=20ng, 40ng or 80ng (24 well plate, 12 well plate or 6 well plate respectively). For experiments including chANP32A, pCAGGS expression plasmid encoding FLAG-tagged chANP32A was included in the transfection mix at a ratio of 4. For replication assays WT PB1 plasmid was pre-expressed, whilst for cRNP stabilisation assays catalytically dead polymerase (PB1 D446Y) was pre-expressed. 20 hours post transfection, cells were infected as described above, with the addition of actinomycin D (5 μg/mL), cycloheximide (100 μg/mL) or DMSO control as indicated. At the appropriate time point, cells were processed for RT-qPCR analysis, imaging analysis or immunoblot as described below.

### Tagged RT-qPCR against vRNA, cRNA and mRNA

For RT-qPCR analysis, 293T or eHAP cells were cultured in 24 well plates, with each condition in triplicate. Following infection/transfection, cells were lysed using buffer RLT or RLT plus (Qiagen), frozen at −80, then total RNA was extracted using either the RNeasy RNA extraction kit (Qiagen) with 30 minute on column DNAseI digest, or the QIAsymphony RNA kit (Qiagen). Quantification for segment 4 and segment 6 vRNA, cRNA and mRNA was based on the tagged primer approach developed by Kawakami et al (24). For each sample, four reverse transcription reactions were set up using 200ng RNA/reaction, RevertAid H Minus Reverse Transcriptase (Thermo Scientific) as per the manufacturer’s instructions, plus a tagged primer targeting either vRNA or cRNA, a tagged polydT (for viral mRNA) or an untagged polydT (for GAPDH internal control). For NA vRNA, cRNA and mRNA, primers used were <u>GGCCGTCATGGTGGCGAAT</u>GAAACCATAAAAAGTTGGAGGAAG, <u>GCTAGCTTCAGCTAGGCATC</u>AGTAGAAACAAGGAGTTT and <u>CCAGATCGTTCGAGTCG</u>TTTTTTTTTTTTTTTTTT respectively, tags underlined)) whilst primers used for HA vRNA, cRNA and mRNA were <u>GGCCGTCATGGTGGCGAAT</u>GGAGAGTGCCCAAAATACGT, <u>GCTAGCTTCAGCTAGGCATC</u>AGTAGAAACAAGGGTGTT and CCAGATCGTTCGAGTCGTTTTTTTTTTTTTTTTTT respectively, tags underlined). Tagged cDNA was then diluted 1 in 10 and quantified using real-time quantitative PCR using Fast SYBR green master mix (Thermo Scientific). Primer pairs used were: CCTTCCCCTTTTCGATCTTG/ GGCCGTCATGGTGGCGAAT (NA vRNA), CTTTTTGTGGCGTGAATAGTG/ GCTAGCTTCAGCTAGGCATC (NA cRNA), CTTTTTGTGGCGTGAATAGTG/ CCAGATCGTTCGAGTCGT (NA mRNA), CATACCATCCATCTATCATTCC/ GGCCGTCATGGTGGCGAAT (HA vRNA), GGGGGCAATCAGTTTCTG/ GCTAGCTTCAGCTAGGCATC (HA cRNA), GATTCTGGCGATCTACTCAACTGTC/ CCAGATCGTTCGAGTCG (HA mRNA) and AATCCCATCACCATCTTCCA/ TGGACTCCACGACGTACTCA (GAPDH). qPCR analysis was carried out in duplicate or triplicate on a Viia 7 real-time PCR system (Thermo Fisher). Fold changes in gene expression relative to either input (0hpi) or mock infected controls (as indicated in figure legends) were calculated using the 2^- ΔΔCT^ method with GAPDH expression as internal control.

### RNAscope/immunofluorescence co-staining

For imaging analysis, cells were cultured on glass cover slips coated in poly-L-lysine in 12 well plates. At the appropriate timepoint, infected cells were washed in phosphate buffered saline (PBS, Gibco) and fixed in 4% paraformaldehyde for 30 minutes, prior to further washes in PBS and dehydration in an ethanol gradient (50% EtOH, 5 mins, 70% EtOH, 5 mins, 100% EtOH, 5 mins, fresh 100% EtOH, 10 mins). Cover slips were stored in 100% ethanol at −20°C until further processing. For RNAscope/immunofluorescence co-staining, RNA was stained first using RNAscope probes (ACDBio). Probes were designed to target PR8 NA vRNA (channel 1) and PR8 NA cRNA/mRNA (+RNA) (channel 2). Cover slips were rehydrated in an ethanol gradient (70% EtOH, 2 mins, 50% EtOH, 2 mins, PBS, 10 mins), treated with protease III diluted 1 in 15 in PBS and staining undertaken using the fluorescent multiplex kit v1, following the manufacturer’s instructions up until and including incubation in the final fluorophore mixture (Fl-Amp4). At this point, cover slips were blocked in PBS with 2% bovine serum albumin (BSA) and 0.1% tween for 30 minutes, incubated in rabbit α-PB2 (catalog no. GTX125926, Genetex) antibody for 1 hour at room temperature (RT), followed by goat α-rabbit AF647 (Invitrogen) plus DAPI for 1 hour, RT. Images were obtained using a Leica SP5 inverted confocal microscope and processing undertaken using FIJI software (33, 34).

### Immunoblot analysis

To confirm equivalent protein expression during replication/cRNP stabilisation assays, cells transfected in parallel were lysed with homemade radioimmunoprecipitation assay (RIPA) buffer (150 mM NaCl, 1% NP-40, 0.5% sodium deoxycholate, 0.1% SDS, 50 mM Tris, pH 7.4) supplemented with an EDTA-free protease inhibitor cocktail tablet (Roche). Lysates were clarified, then mixed with 4x Laemmli sample buffer (Bio-Rad) with 10% β-mercaptoethanol. Membranes were probed with rabbit α-vinculin (EPR8185, Abcam) or mouse α-tubulin (ab7291, Abcam) and mouse α-NP (C43, Abcam), followed by near-infrared secondary antibodies (IRDye 680RD goat anti-rabbit (IgG) secondary antibody (Abcam) and IRDye 800CW goat anti-mouse (IgG) secondary antibody (Abcam)). Western blots were visualised using an Odyssey imaging system (Li-Cor Biosciences).

### Statistical analysis

Statistics throughout this study were performed using one-way analysis of variance (ANOVA) or student’s T-test as described in figure legends.

We thank Ervin Fodor and Ecco Staller for the kind gift of the ANP32A/B/E triple knockout eHAP cells. We thank Andreas Bruckbauer and Stephen Rothery at the Facility for Imaging by Light Microscopy (FILM) at Imperial for support with microscopy.

This work was supported by Wellcome Trust grant 205100. In addition, O.C.S. was supported by a Wellcome Trust studentship, A.B.R. was supported by a Medical Research Council (MRC) studentship, and T.P.P. was supported by Biotechnology and Biological Sciences Research Council (BBSRC) grant BB/R013071/1.

O.C.S., A.M.B., C.M.S and W.S.B. conceived and planned experiments; O.C.S. and A.M.B. performed experiments and analysed data, O.C.S., T.P.P., C.M.S. and W.S.B. provided supervision, O.C.S. and W.S.B. wrote the original manuscript, O.C.S., A.M.B., T.P.P., C.M.S. and W.S.B. reviewed and edited the manuscript.

The authors declare no competing interests.

---

**Figure S1.**
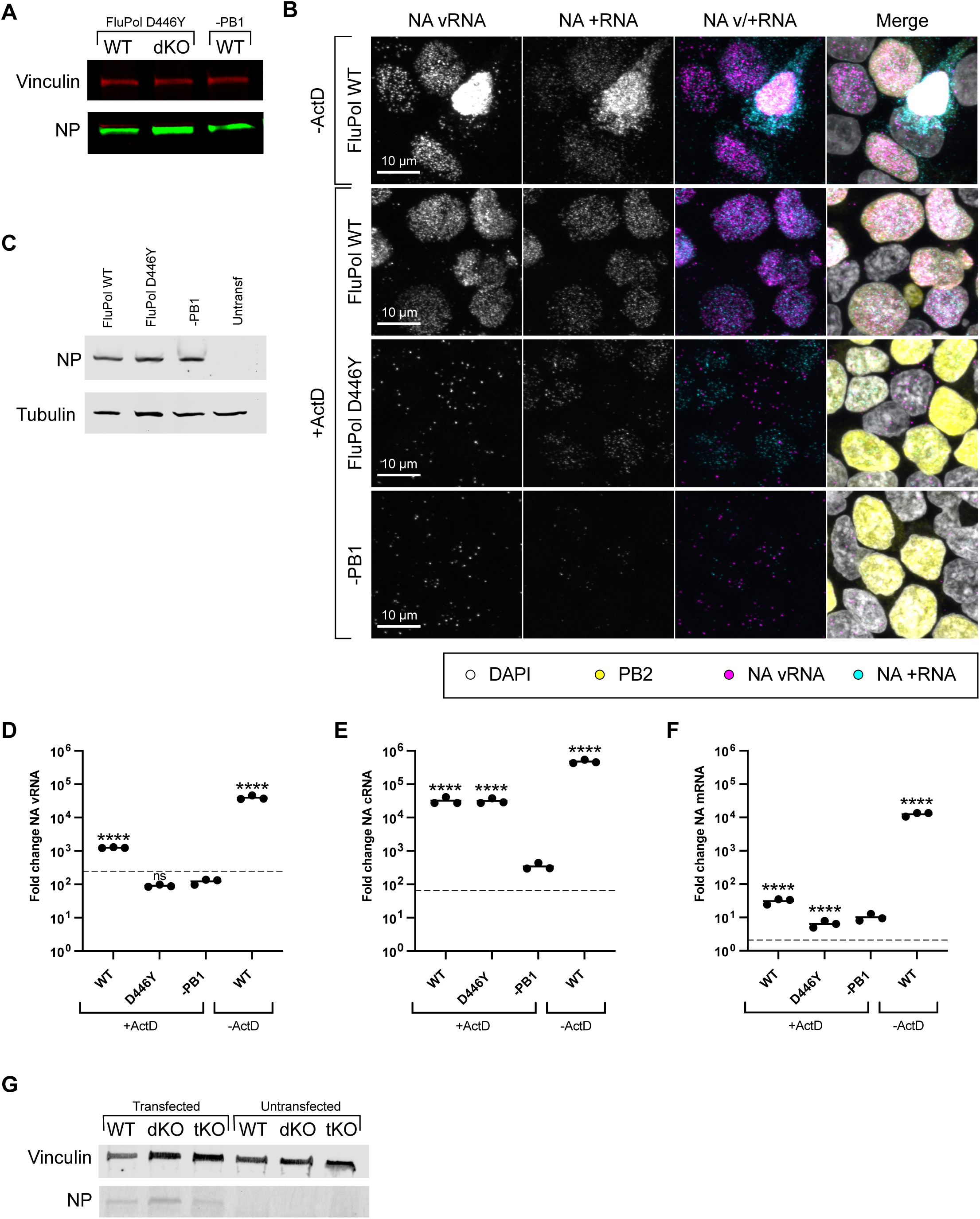
Supporting data for figure 2. (A) Matched western blot for Fig. 2B,C,D. (B-F) Validation of replication and stabilisation assays with ActD analysed by RNA FISH. Assays were performed in HEK 293T cells, following infection with PR8 (MOI=3). (B) Accumulation of NA vRNA and +RNA, 3hpi, analysed by RNAscope with indirect immunofluorescence against PB2. Pre-transfected PR8 polymerase mixes are indicated on the left-hand side: FluPol WT=PB2/PB1/PA/NP; FluPol D446Y=PB2/PB1 D446Y/PA/NP; -PB1=PB2/PA/NP. Images are representative maximum intensity projections. (C) Matched western blot confirming equal transfection of different polymerase mixtures. (D,E,F) Matched RT-qPCR analysis of segment 6 (D) vRNA (E) cRNA and (F) mRNA accumulation, 6hpi. Pre-transfected polymerase mixes are indicated on the x-axis. Dotted line indicates input levels of RNA (0hpi). Fold change was calculated over mock infected cells. n=3 biological replicates, plotted as mean ± s.d.. Significance compared to –PB1 was assessed using one-way ANOVA with Dunnett’s multiple comparison test, following log transformation. (G) Matched western blot for Fig. 2G,H,I. Untransf= Untransfected; hpi=hours post infection. ns=not significant; ****=p<0.0001.

**Figure S2.**
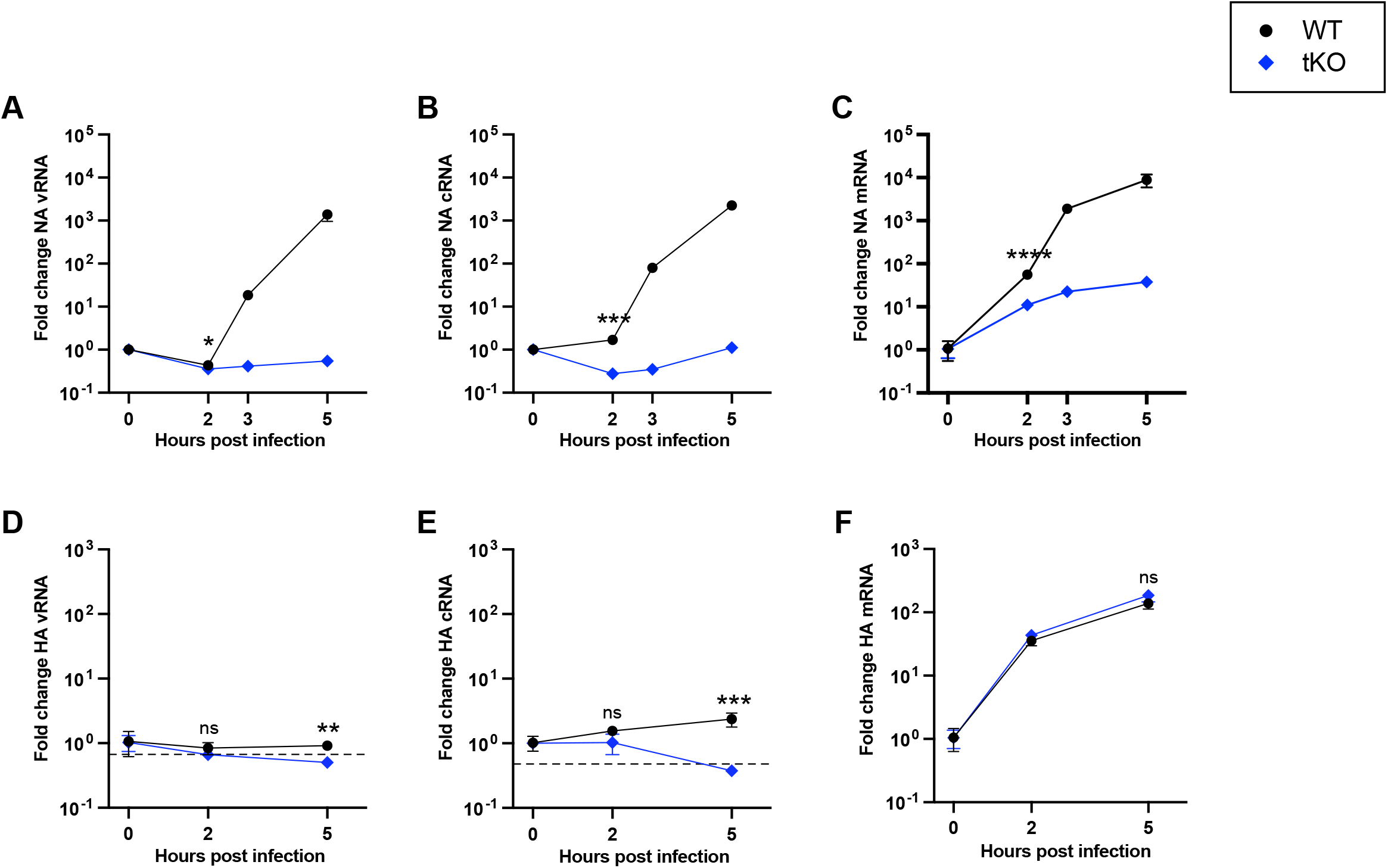
Human ANP32 proteins play a direct role in primary cRNA synthesis of H5N1 5092. (A-C) Segment 6 (A) vRNA (B) cRNA or (C) mRNA accumulation over time in eHAP WT vs tKO cells following infection with 5092K (MOI=3). Fold change was calculated over input (0hpi). n=3 biological replicates, plotted as mean ± s.d.. Significance at 2hpi was assessed using an unpaired t-test following log transformation. (D,E,F) cRNP stabilisation assay with CHX in eHAP WT and tKO cells. Segment 4 (D) vRNA (E) cRNA or (F) mRNA accumulation following infection with 5092K (MOI=3). Fold change was calculated over input (0hpi). Dotted line indicates background levels of vRNA/cRNA present in WT cells transfected with a control plasmid mix minus PB1, 5hpi. n=3 biological replicates, plotted as mean ± s.d.. Significance was assessed using multiple unpaired t-tests following log transformation, corrected for multiple comparisons using false discovery rate. hpi=hours post infection. ns=not significant; *= p<0.05; **=p<0.01; ***=p<0.001; ****=p<0.0001. e 1. cRNA synthesis is inhibited in human cells lacking ANP32A/B. (A-C) Segment 4 (A) vRNA RNA or (C) mRNA accumulation over time in eHAP WT vs dKO cells following infection with PR8 =3). Fold change was calculated over input (0hpi). n=3 biological replicates, plotted as mean ± Significance was assessed at 2hpi using an unpaired t-test following log transformation. (D-F) ent 6 (D) vRNA (E) cRNA or (F) mRNA accumulation over time in eHAP WT vs dKO cells follow-fection with PR8 (MOI=3). Fold change was calculated over mock infected cells. n=3 biological ates, plotted as mean ± s.d.. Significance was assessed using multiple unpaired t-tests following ransformation, corrected for multiple comparisons using false discovery rate. hpi=hours post tion. ns=not significant; **=p<0.01; ***=p<0.001; ****=p<0.0001.

